# 10 years of EPOC: A scoping review of Emotiv’s portable EEG device

**DOI:** 10.1101/2020.07.14.202085

**Authors:** Nikolas S Williams, Genevieve M McArthur, Nicholas A Badcock

## Abstract

**BACKGROUND:** Commercially-made low-cost electroencephalography (EEG) devices have become increasingly available over the last decade. One of these devices, Emotiv EPOC, is currently used in a wide variety of settings, including brain-computer interface (BCI) and cognitive neuroscience research.

**PURPOSE:** The aim of this study was to chart peer-reviewed reports of Emotiv EPOC projects to provide an informed summary on the use of this device for scientific purposes.

**METHODS:** We followed a five-stage methodological framework for a scoping review that included a systematic search using the Preferred Reporting Items for Systematic Reviews and Meta-Analyses Extension for Scoping Reviews (PRISMA-ScR) guidelines. We searched the following electronic databases: PsychINFO, MEDLINE, Embase, Web of Science, and IEEE Xplore. We charted study data according to application (BCI, clinical, signal processing, experimental research, and validation) and location of use (as indexed by the first author’s address).

**RESULTS:** We identified 382 relevant studies. The top five publishing countries were the United States (n = 35), India (n = 25), China (n = 20), Poland (n = 17), and Pakistan (n = 17). The top five publishing cities were Islamabad (n = 11), Singapore (n = 10), Cairo, Sydney, and Bandung (n = 7 each). Most of these studies used Emotiv EPOC for BCI purposes (n = 277), followed by experimental research (n = 51). Thirty-one studies were aimed at validating EPOC as an EEG device and a handful of studies used EPOC for improving EEG signal processing (n = 12) or for clinical purposes (n = 11).

**CONCLUSIONS:** In its first 10 years, Emotiv EPOC has been used around the world in diverse applications, from control of robotic limbs and wheelchairs to user authentication in security systems to identification of emotional states. Given the widespread use and breadth of applications, it is clear that researchers are embracing this technology.

## Introduction

Electroencephalography (EEG) is a continuous recording of the electrical activity generated by groups of neurons firing in the brain. An EEG typically comprises recordings of activity present at multiple sites on the head, indexed using metal electrodes placed on the scalp. EEG recordings can be inspected by sight for signs of brain dysfunction (e.g., epilepsy), or can be processed to produce spectral analyses of the electrical activity over a period of time, and event-related potentials (ERPs) that reflect the average pattern of electrical activity generated by a particular stimulus (e.g., a speech sound, a face, a written word).

EEG is one of the oldest neuroscientific techniques in use today. Since the first human recordings published by Hans Berger in 1929 [see 1, for a history], EEG has become a popular tool for neuroscientists due to its non-invasive nature and high temporal resolution. The technique has matured over the decades due to advances in technology, which has allowed for greater instrument sensitivity and better signal processing techniques. What used to be analogue signals scribed onto rolls of paper are now digital recordings stored on hard drives, ready for processing using a myriad statistical and mathematical techniques.

In recent years, one of the biggest evolutions in EEG applications has been the development of consumer-grade devices. Not only do these devices make acquiring EEG signals easier, but they can do so in natural environments outside the traditional laboratory setting. In 2009, a biotech company, Emotiv Systems, released EPOC, a consumer-oriented EEG device. EPOC was originally designed and marketed as a hands-free videogame device, placing it within the class of brain-computer interface (BCI) devices. As one of the first EEG devices available to consumers, EPOC’s release demonstrated the feasibility of low-cost neuroimaging outside of research laboratories. The next 10 years saw the EPOC developer re-established as Emotiv Inc., a second iteration of the device called EPOC+, and the market of EPOC evolve to include research applications. Neuroscientists, keen to take advantage of efficiency increases and budget decreases, saw an opportunity in EPOC for user-friendly research at a fraction of the cost of traditional research-grade EEG systems.

In the decade since its release, EPOC has been used in hundreds of scientific applications as its ease of setup and low price-point make it an appealing option for researchers and engineers. The first published works using EPOC appeared in 2010, describing the use of EPOC in BCI applications [2-4]. In 2011, the first study using for experimental research was published [5]. Two years later, studies validating the use of EPOC in experimental research began to emerge [6-8]. In the years that followed, EPOC appeared in many conference proceedings and journal articles, suggesting its wide adoption as an EEG device. In our laboratory, we have successfully converted the EPOC into an ERP device, which we have validated against a research-grade system [8-10]. In addition, our department has integrated EPOC into the Bachelor of Cognitive and Brain Sciences as a practical demonstration of neuroscience methodology [11].

Given the demonstrated validity of the EPOC as a research tool, as well as its low cost, researchers around the world are understandably curious about what the EPOC system can and cannot be used for. This has inspired a number of reviews of EPOC’s use in specific domains such as BCI [12-17], cognitive enhancement [18, 19], stress detection [20], and education [21]. However, no review has considered the use of the EPOC across multiple domains. In addition, while other reviews have focused on portable EEG devices in general [22-26], none have focused on the EPOC device specifically.

With this gap in knowledge in mind, we aimed to carry out a scoping review of studies that have used the EPOC as an EEG and ERP device to understand the use and location of EPOC research to date. We followed the framework put forth by Arksey and O’Malley (27 p. 21) for conducting a scoping review, where, in contrast to a systematic review, a scoping review does not seek to answer a narrowly-defined research question but to examine and describe the “extent, range, and nature of research activity”. We followed the five stages described by Arksey and O’Malley (27), which were:

Stage 1: identifying the research question.

Stage 2: identifying relevant studies.

Stage 3: study selection.

Stage 4: charting the data.

Stage 5: collating, summarising, and reporting the results.

Additionally, we followed the Preferred Reporting Items for Systematic Reviews and Meta-analyses Extension for Scoping Reviews (PRISMA-ScR) guidelines [28]. See supporting information (S1 PRISMA-ScR) for the checklist.

### Stage 1: Identifying the Research Question

We sought to answer the question of where (i.e., locations) and how (i.e., applications) EPOC has been used in research settings. In addressing this question, we aim to facilitate decision-making about EPOC useability and expect this review may be particularly beneficial for researchers who are searching for inexpensive neuroscience techniques. It may also be useful for clinicians in the development of BCI-assisted technologies that support people with physical limitations.

## Emotiv EPOC

There have been two versions of Emotiv’s device, EPOC and EPOC+. The primary difference is that EPOC+ can capture data at 128 Hz and 256 Hz sampling rates whereas EPOC samples at 128 Hz only. We reviewed projects using both devices in this scoping review, but for simplicity we will refer to both versions as EPOC.

### Stage 2: Identifying Relevant Studies

The first author conducted a systematic search of the literature by retrieving records from the following online bibliographic databases: (a) PsychINFO, (b) MEDLINE, (c) Embase, (d) Web of Science, and (e) IEEE Xplore. These widely-used databases cover a large breadth of fields, including psychology, cognitive science, medicine, and engineering. Searches included peer-reviewed studies conducted with human participants and written in English. Searches included studies published from 2009 onwards (i.e., the year EPOC was released). To find records in each database, we used the following search strings in conjunction with wildcards to capture keyword variations: Emotiv, EPOC, electroencephalograph, EEG, event-related, ERP. For example, in PsychINFO we used: (Emotiv OR EPOC) AND (electroenceph* OR EEG OR event-related OR event related OR ERP). The initial search was conducted in June of 2018. A second search was conducted in February of 2019 and a third search was conducted in February 2020.

Fig 1 outlines the Preferred Reporting Items for Systematic Reviews and Meta-analyses (PRISMA) flowchart for this review (Moher, Liberati, Tetzlaff, & Altman, 2009). In brief, we identified 724 articles via the database search. This included 249 duplicate articles resulting in 475 articles after removal.

**Fig 1.**
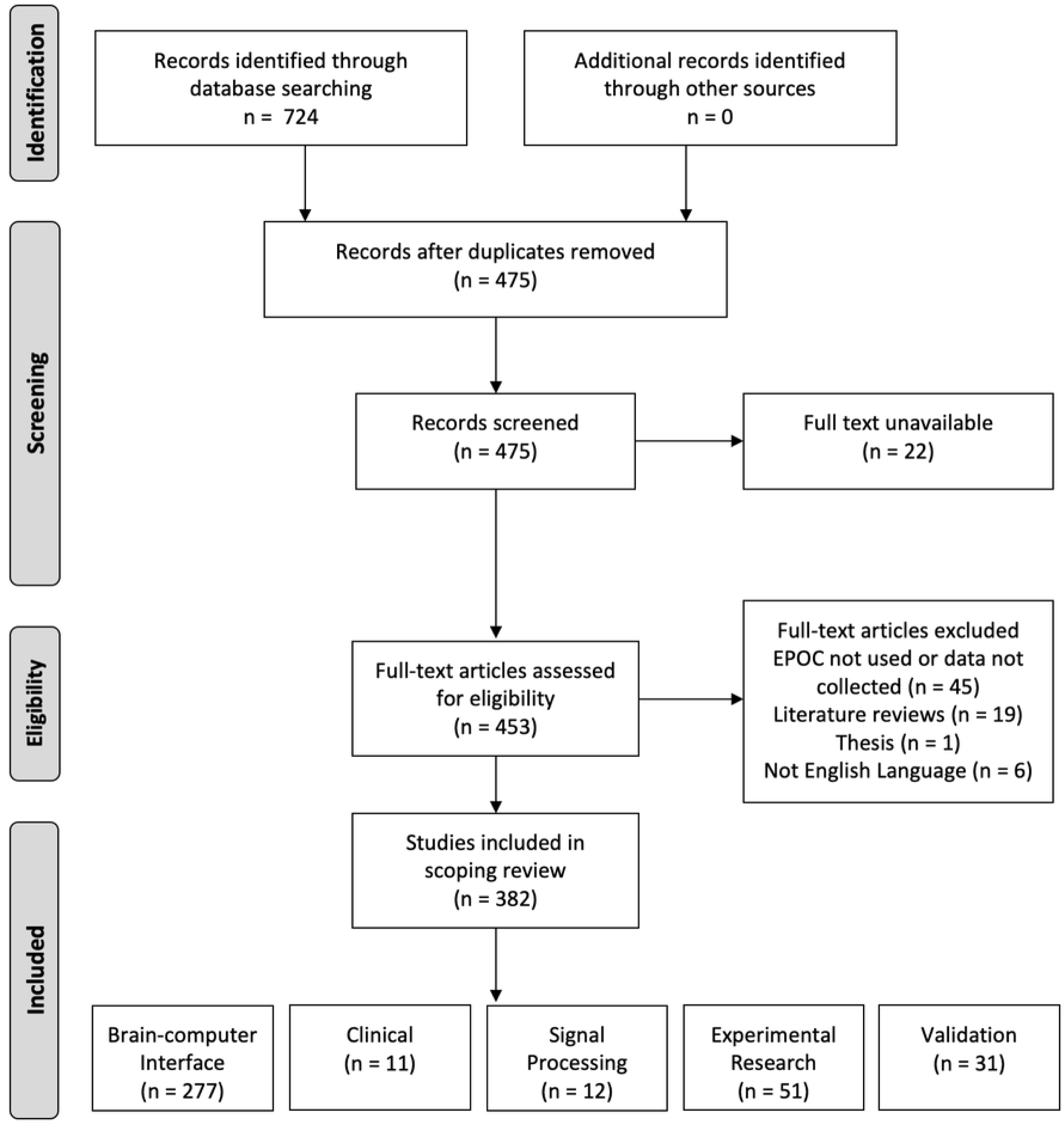
Preferred Reporting Items for Systematic Reviews and Meta-analyses (PRISMA) flowchart.

### Stage 3: Study Selection

We excluded twenty-two records for which the full-text could not be retrieved and screened the remaining articles according to the following eligibility criteria: (1) EPOC device used; (2) Actual data collected; (3) Articles published in peer-reviewed journals or conference proceedings. We removed seventy-one studies that failed at least one of these criteria. These included publications in which EPOC appeared as the acronym for *Effective Practice and Organisation of Care*, in which no actual EEG data was collected, which were not written in English, or were literature reviews.

The final number of studies included in this review was 382. Of these, 252 were conference proceedings and 130 were journal articles. As the conference proceedings in this review meet the criteria of peer review, we did not distinguish between conference proceedings and journal articles. However, Fig 2 provides a breakdown of the types of studies over the years included in this review.

**Fig 2.**
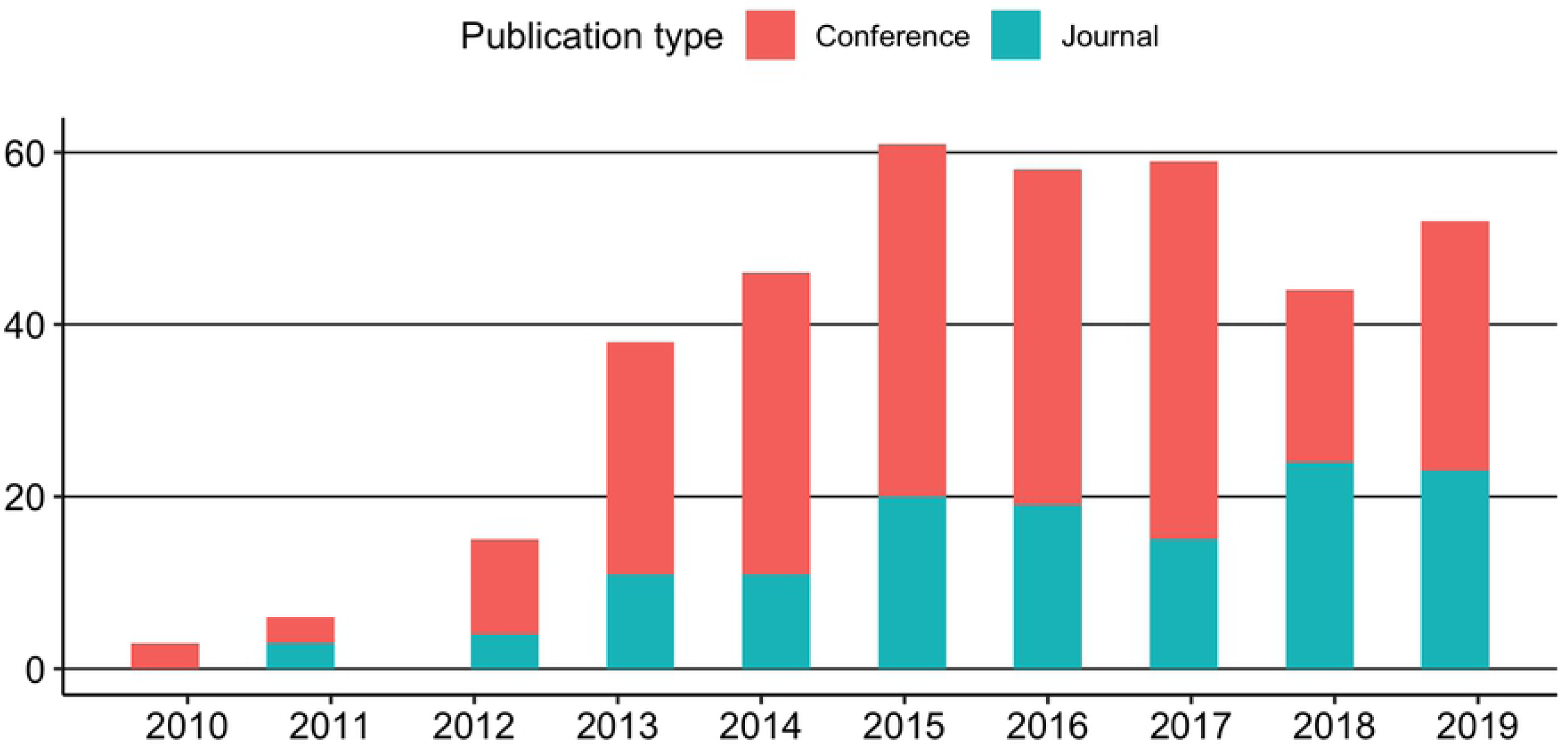
Number of EPOC studies by type from 2010 to 2019.

### Stage 4: Charting the data

We charted the data by recording relevant information from each record. This information included the author(s), year of publication, study location, and aims of the study. We classified each study according to its aim into one of five categories: (1) EPOC used as a BCI device (e.g., control of a wheel chair); (2) EPOC used as a clinical tool (e.g., to assess depression); (3) EPOC used to collect EEG data for developing or refining EEG signal processing techniques (e.g., to reduce artefacts in EEG data streams); (4) EPOC used as a theory development tool (e.g., to examine EEG signatures in cognitive tasks); and (5) studies aimed at validating EPOC as an EEG device (e.g., comparing EPOC to research-grade EEG systems). See Table 1 for descriptions and number of studies assigned to each category.

**Table 1.**
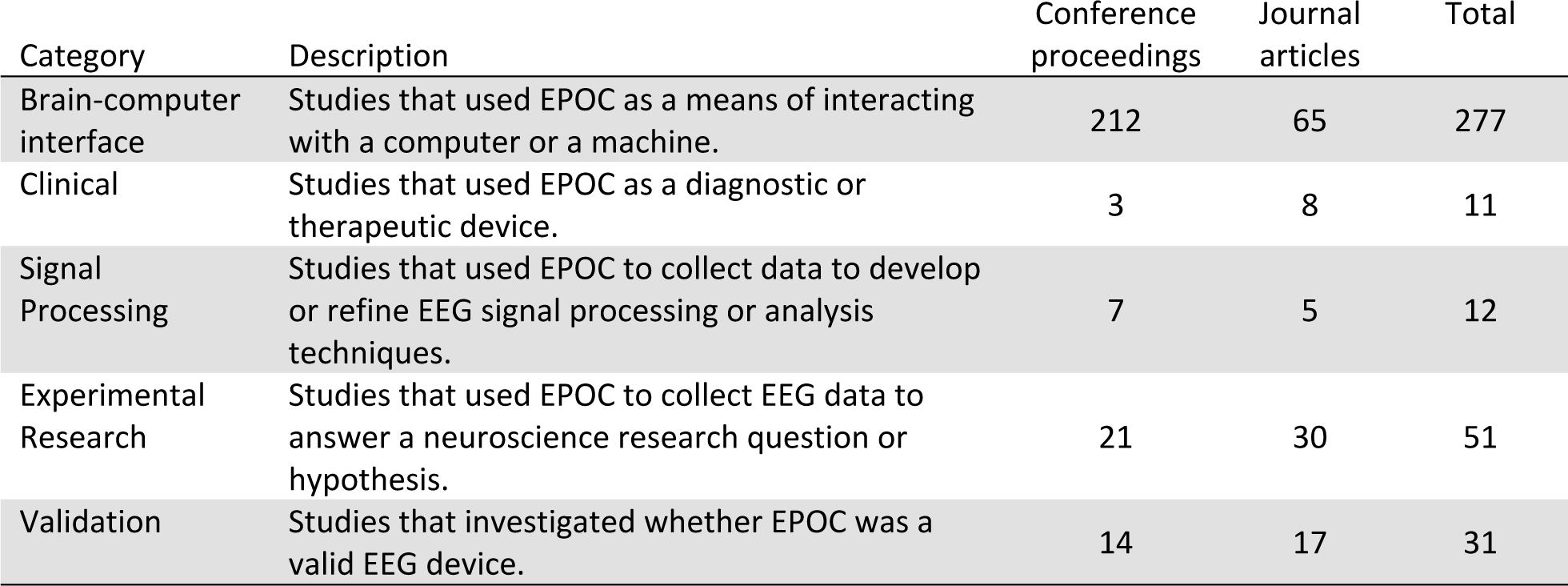
Category descriptions and counts of EPOC-related studies.

To chart the location of EPOC studies, we used the first authors’ corresponding address. To visualise the global distribution of EPOC studies, we obtained latitude and longitude coordinates from a world cities database (https://simplemaps.com/data/world-cities). If cities did not have coordinate information in the database, we performed a Google search and entered the coordinates manually. See supporting information (S2 Appendix) for data extracted and charted.

### Stage 5: Collating, Summarising, and Reporting the Results

The first three EPOC-related studies were published in the year after its release, 2010. These initial studies were all related to BCI: two P300 classification studies [2, 4] and a robotic arm control study [3]. A year later, 2011, saw the first study published using EPOC for experimental research [5]. This study examined the relationship between EEG, personality, and mood on perceived engagement. The first publications aimed at validating EPOC appeared in 2013 [6-8, 29, 30].

Overall, the number of EPOC studies showed a steady increase from 2010 (n = 3) to 2015 (n = 61), after which the numbers fell to 58, 59 and 44 in 2016, 2017, and 2018 respectively, and then increased to 52 publications in 2019 (see Fig 3). While the true reason for this pattern is unknown, it may well reflect a change in the licensing of Emotiv software, which switched to a subscription-based license in 2016 (previously the license involved a one-time fee). This increased the cost of the EPOC for research, which may explain the declining in publications in 2015 – 2018. The resurgence in 2019 could be acceptance of the licensing fee as the new standard and being factored into budgets and grant applications. It remains to be seen how this fee structure will impact EPOC use in the future.

**Fig 3.**
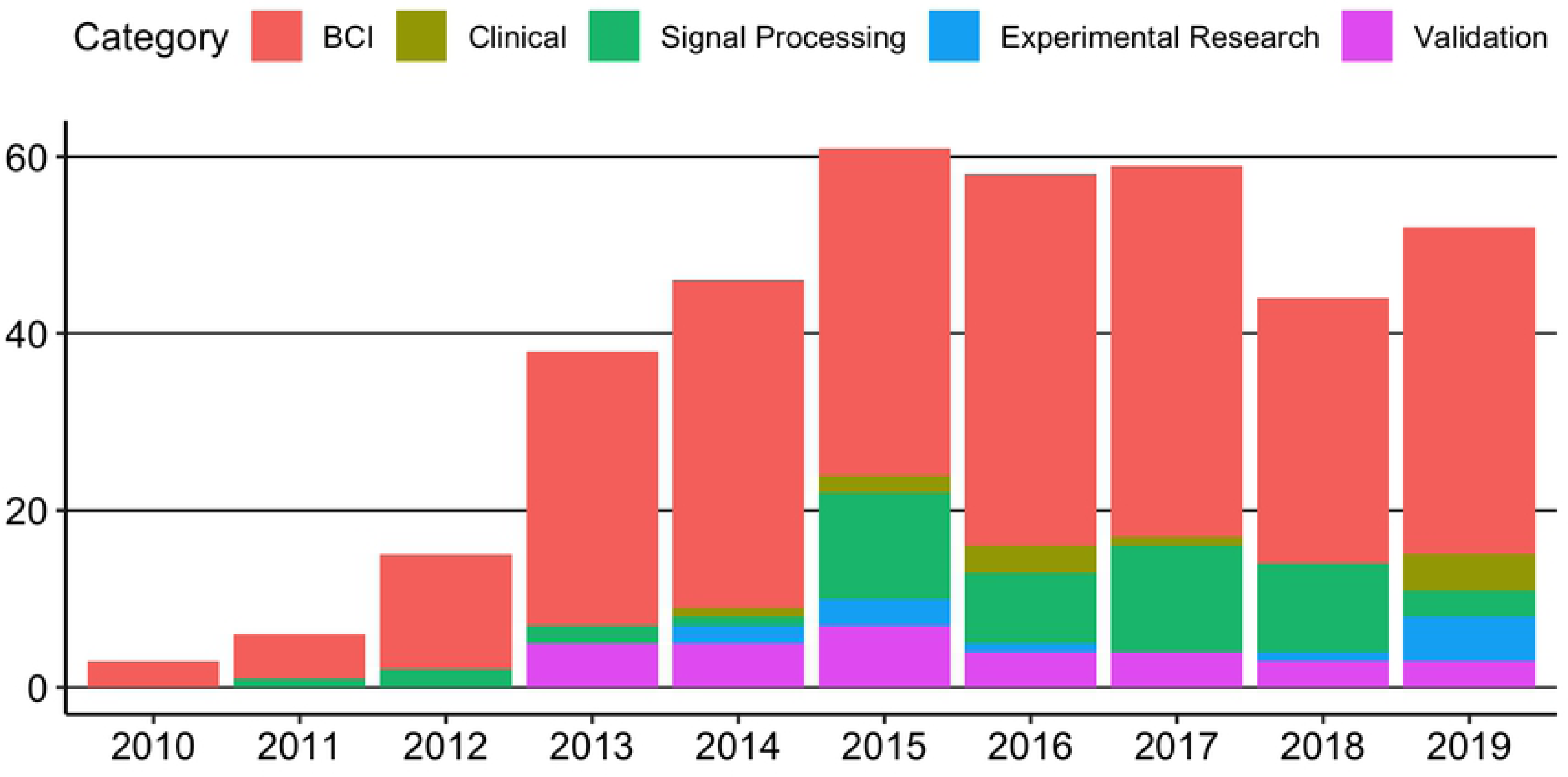
Number of EPOC studies from 2010 to 2019 by study application.

## Location

In the years 2009 to 2019, the five countries that published the most EPOC studies were the United States (n = 35), India (n = 25), China (n = 20), Poland (n = 17), and Pakistan (n = 17). The five individual cities that published the most EPOC studies were Islamabad (n = 11), Singapore (n = 10), Bandung, Indonesia (n = 7), Cairo (n = 7), and Sydney (n = 7). See Fig 4 for overall global distribution of studies covered by this systematic review.

**Fig 4.**
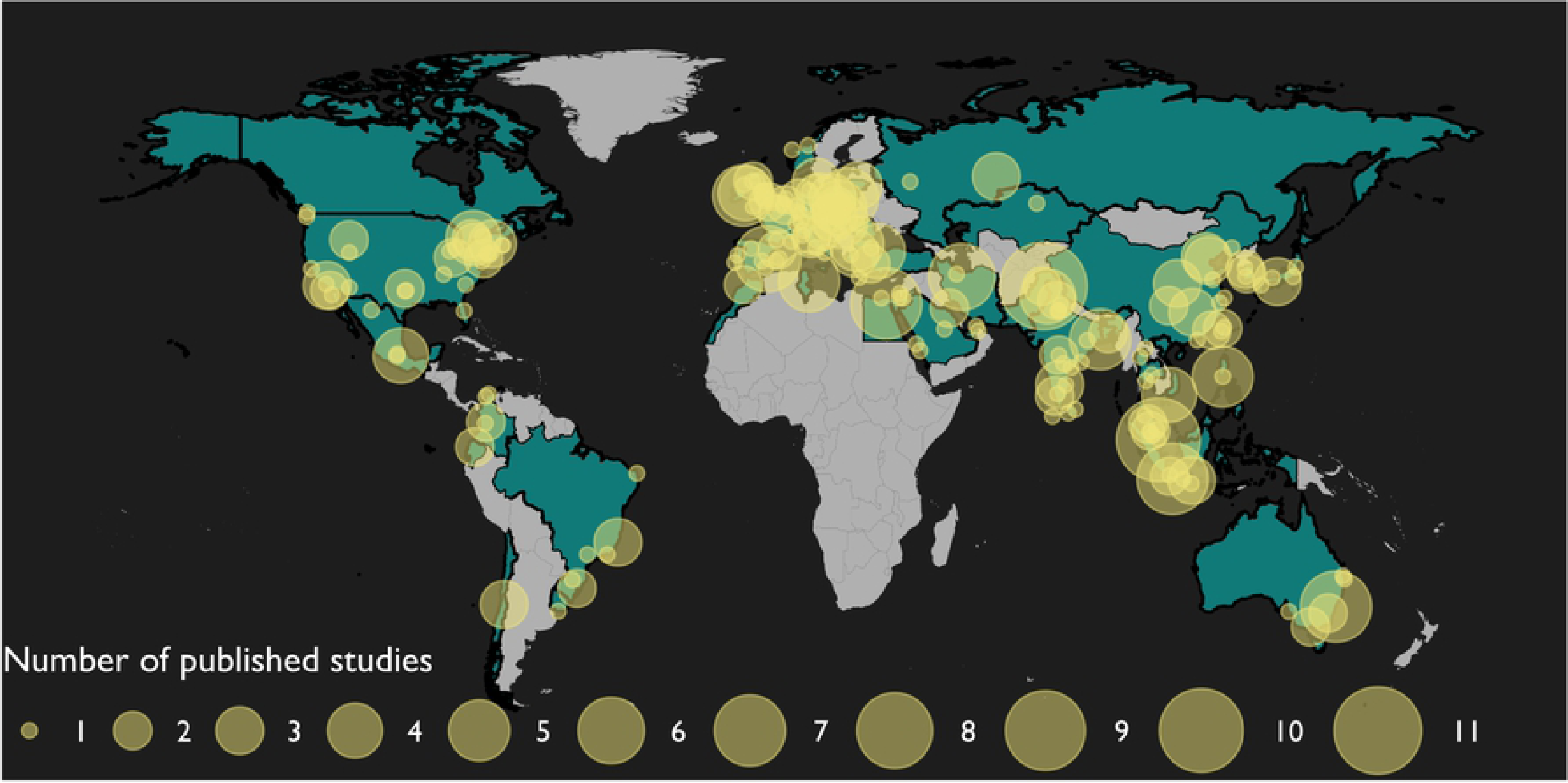
Global distribution of EPOC studies from 2010 to 2019.

## Applications

### BCI

BCI applications represented the majority of EPOC studies (approximately 73% of studies in this review). To better characterise BCI studies, we further classified them into four subcategories: (a) biometrics, (b) device control, and (c) state recognition, and (c) general classification. See Table 2 for description of each subcategory.

**Table 2.** Descriptions and study counts of EPOC-related BCI subcategories.

| BCI Subcategory | Description | Study count |
| --- | --- | --- |
| Biometrics | Studies that used EPOC to characterise EEG signatures for identification/authentication of individual users. | 11 |
| Device control | Studies that used EPOC to enable control of devices in an individual's environment (e.g., wheelchairs, appliances). | 97 |
| State recognition | Studies that used EPOC to recognise human physical, mental, or affective states. | 94 |
| General classification | Studies that used EPOC to collect EEG data for the classification of EEG signatures, without explicit applications associated with other BCI subcategories in this table. | 75 |

### BCI biometrics

Much like a fingerprint or a password, individual brain signatures can identify individuals, granting them access to systems or facilities. As the variation between individuals’ brain waves can be quite complex, the use of individual EEG signatures as a biometric indicator represents a promising application of portable EEG technology. A total of 11 studies used EPOC to investigate EEG in the context of user-authentication and security. The earliest biometric EPOC study was published in 2013 and used a P300 speller paradigm to investigate the feasibility of using EEG classification for user authentication [31]. Recently, study designs and classification methods have grown more sophisticated. For example, Moctezuma, Torres-Garcia (32) used feature extraction and classification to distinguish between individuals’ EEG signatures while they imagined speaking words. Likewise, Seha and Hatzinakos (33) also employed feature extraction, in this case on auditory evoked potentials, to accurately (> 95%) discriminate between individuals. Compared to BCI studies in general, relatively few EPOC studies have focused on biometry. Nevertheless, there has been a general increase in biometric studies since 2013 and this field represents one of the many practical applications of portable EEG technology.

### BCI device control

The control of external devices, such as prostheses or wheelchairs, is the most straightforward application of EEG-based BCIs. A total of 97 studies, representing 35% of BCI studies and 25% of studies overall, used EPOC as a means of controlling or interacting with machines in users’ environment. P300 spellers are perhaps the most well-known type of BCI interface that fall under this category. P300 speller interfaces exploit the well-documented and robust signature observed as deflection in an ERP waveform in a response to a target stimulus. By capitalising on the P300, a computer can detect when a target letter flashes on a screen thereby allowing selection of letters without physical interaction. Though there were several EPOC studies in this review that investigated traditional P300 speller BCI interfaces [34-37], others harnessed P300 for such purposes as interacting with navigation systems [38] and robotic devices [39]. Suhas, Dhal (40) investigated using ERPs to control a light bulb and a fan, with an eye towards giving physically disabled individuals control of ‘smart’ appliances. Other studies have employed EPOC as a means of controlling robots [41-47], tractors [48], and drones [49]. Practical and effective BCI device control using EEG has the potential to benefit a large population, such as individuals who have lost the use of motor functions. For this reason, this area of research has received much attention and it should be expected to continue to do so.

### BCI state recognition

Characterising and identifying cognitive or affective states using EEG is critical for many BCI paradigms and is a hallmark of neurofeedback applications. Many researchers have used EPOC to achieve this. For example, an early EPOC study attempted to recognise EEG patterns when participants imagined pictures [50]. More recent studies have used sophisticated algorithms to identify cognitive states, such as confusion [51], fatigue [52] and emotions [53-55]. Identifying an individual’s mental state can help to improve human performance in demanding situations. For example, Binias, Myszor (56) used an EPOC to develop algorithms aimed at helping pilots respond more quickly to unanticipated events. Studies like these demonstrate that the rapid and accurate identification of cognitive and affective states, even before conscious recognition, may lead to safer roads and skies.

### BCI general classification

A total of 75 BCI studies were not readily classifiable in the above subcategories as they were not concerned with a direct application of research. Rather, these studies aimed to increase the usability of BCI technology through the development and refinement of EEG classification algorithms. For example, Perez-Vidal, Garcia-Beltran (57) collected EEG data with EPOC in order to determine the effectiveness of a machine-learning algorithm for correctly identifying P300 evoked potentials. In this example, direct use of the P300 was not directly used for interfacing with a specific device/machine. Rather, the central focus was on the algorithm itself. We categorised these types of studies as general classification studies.

### Clinical

The small form factor and ease of setup make portable EEG devices ideal for use in clinical settings in which the objective is to treat or diagnosis health conditions. Eleven studies in this review used EPOC for this purpose, with six studies aimed at using EPOC specifically for a therapeutic purpose. For example, studies have used EPOC to provide neurofeedback for motor rehabilitation [58, 59] or for the treatment of depression [60] and pain [61, 62]. Five studies used EPOC as a diagnostic tool with the aims of assessing conditions such as depression [63], attention deficit hyperactivity disorder [64, 65], or encephalopathy [66]. Yet another study used EPOC to monitor changes in the nervous system of a group of Turkish researchers who visited Antarctica [67].

### Signal processing

Twelve studies in this review aimed to improve EEG signal processing techniques used with EPOC data. For example, Sinha, Chatterjee (68), Soumya, Zia Ur Rahman (69), and Jun Hou, Mustafa (70) used EPOC to test techniques aimed at reducing EEG artefacts and noise. Additionally, Moran and Soriano (71) compared different techniques for maximising EPOC EEG signal quality while Petrov, Stamenova (72) and Shahzadi, Anwar (73) investigated remote EEG transmission and processing. These studies are important as the signal-to-noise ratio of EEG can be small and techniques aimed at increasing it can broaden the utility of EEG devices. In addition, the increasingly distanced nature of research and clinical diagnostics necessitates the development of effective data transmission pipelines.

### Experimental Research

We identified a total of 51 experimental research studies that used EPOC incidentally to answer questions related to brain function. That is, researchers could have used any EEG device to collect data but they chose EPOC. Most of these studies were directly concerned with investigating EEG signatures related to certain processes, situations, or tasks. Many were cognitive in nature including EEG signatures related to cognitive load [74-77], alertness [78], distraction [79], learning styles [76], semantic association [80-82], and memory [83]. Other studies examined EEG signatures related to perception. These included spatial perception [84], taste perception [85], olfactory perception [86], and visual perception [87, 88].

Studies examining social phenomena also constituted a large proportion of EPOC research projects. For example, we found studies in which EPOC was used to investigate consumer behaviour and preference [89-92]. Other socially-oriented studies examined the EEQ patterns associated with conformity [93], deception [94], perception of quality, [95], and motivation and interest in an educational environment [96].

Researchers also used EPOC to better understand ailments or disorders. These types of studies are contrasted with those in the *clinical* category where publications were aimed at *treating* ailments or disorders, rather than *investigating* the ailments or disorders. For example, Askari, Setarehdan (97), Askari, Setarehdan (98) used the device to investigate neural connectivity in autism. Similarly, Fan, Wade (99) used EPOC to collect EEG data with the aim of building models to accurately identify cognitive and affective states in autistic individuals while driving. In addition to autism, other studies examined the EEG signatures associated with bipolar disorder [100] and mild cognitive impairment [101].

Many research studies were more action-oriented. These types of studies used EPOC to characterise the EEG signatures associated with video games [102, 103], driving [104-106], moving through urban [107] or virtual [108] environments, and performance of specialised tasks [109, 110].

### Validation

Assessing the validity of a device is an important step in establishing its widespread implementation. If an EEG system cannot be demonstrated to accurately capture the data it purports to, then any conclusions drawn from this data are questionable. We identified thirty-one studies that tested the validity of EPOC as a research-grade EEG device. The first EPOC validation studies appeared in 2013. In this year there were five studies assessing the capabilities of EPOC. These studies assessed the accuracy of P300 identification [6], the validity of affect signatures [7], and whether EPOC could be used to collect valid ERPs [8, 29, 30]. Another five EPOC validation studies were published in 2014 before peaking in 2015 (n = 7) and then declining in 2016, 2017, 2018, and 2019 (n = 4, n = 4, n = 3, and n = 3, respectively).

Studies varied in both approach and intended application of the validation. Some did not use a benchmark device with which to compare EPOC. For example, Rodriguez Ortega, Rey Solaz (111) compared EPOC-captured affect signatures to those demonstrated in the literature. Another simply aimed to determine the classification accuracy of EPOC in P300 tasks [112]. However, most studies compared the EPOC to the performance of other research- or consumer-grade EEG devices. Four validation studies compared auditory ERPs between systems [8, 10, 113]. Three studies compared visual ERPs between systems [9, 114 McDowell, & Hairston, 2014, 115, 116]. Tello, Müller (117) also conducted a visual-related validation study in which they compared EEG device performance on steady-state visual evoked potential (SSVEP) tasks, while Szalowski and Picovici (118) tested the capacity of EPOC to distinguish between different SSVEP experimental parameters. Melnik, Legkov (119) compared the performance of multiple systems on both visual ERPs and SSVEPs. Also in the visual domain, Kotowski, Stapor (120) examined the capacity of EPOC to collect ERPs of emotional face processing. Takehara, Kayanuma (121) compared the performance of EPOC to another device on capacity to capture event-related desynchronization while Grummett, Leibbrandt (122) conducted a comprehensive validation study that compared EPOC to other devices on power spectra, ERPs, SSVEPs, and event-related desynchronization/synchronisation.

Some studies validated EPOC’s capacity to measure cognitive indicators with one study comparing devices’ capture of cognitive load signatures [123 Sinharay, & Sinha, 2014], and another comparing the performance of systems during cognitive tasks using time and frequency analyses [124]. Likewise, Naas, Rodrigues (125) tested whether EPOC could enhance cognitive performance in neurofeedback tasks.

Three studies validated EPOC for BCI use by comparing its performance to other device performance on P300 speller tasks [6, 126, 127]. Two others compared the performance of devices on motor imagery tasks [128, 129]. Finally, Maskeliunas, Damasevicius (130) compared the capacity of devices to recognise mental states.

Since 2013, many studies have sought to determine the validity of EPOC. While assessment of the conclusions of these studies are outside the aims of this scoping review, what can be noted is that quantity of studies demonstrates researchers’ interest in employing these devices in their work.

## Limitations

This scoping review has some limitations. With nearly 400 records selected, the charting phase represented an enormous undertaking. Although the review employed a systematic methodology using PRISMA guidelines and searching a broad array of databases, it was impossible to include every study that used EPOC. We deliberately omitted common systematic search strategies, such as grey-literature searching, hand searching, and backward citation searching. We did this as inclusion of these strategies would not have added enough value to justify the additional time and resources. We believe this scoping review represents a quality characterisation of EPOC research and satisfies the stated aims of the project. In addition, like all scientific reviews, its success depends on the search terms. If a publication did not contain ‘Emotiv’ or ‘EPOC’ in the title, abstract, or keywords, then it did not appear in our search. We could have overcome this limitation by broadening our search terms. However, we again believe our search constraints produced an accurate characterisation of the EPOC literature, without creating an unwieldy scoping dataset.

## Conclusion

In this scoping review, we aimed to chart the location and purpose of EPOC-related research. In doing so, we have outlined the many studies that have used Emotiv EPOC as an EEG acquisition device. From BCI applications to experimental research studies to clinical environments, the last 10 years has seen diverse implementation of EPOC. Global use and a low financial barrier likely facilitate research in areas of limited resources. Considering the cost of a research-grade EEG system, it is not hard to imagine scientists and engineers in developing nations embracing EPOC as an ideal device with which to conduct neuroscience research. In addition, this device (and devices like it) may enable collection of data that would be impossible with traditional EEG devices. For example, Parameshwaran and Thiagarajan (131) used an EPOC in both rural and urban settings in India to demonstrate differences in EEG signatures related to factors such as socioeconomic status, exposure to technology, and travel experience.

We expect that this review will provide a useful reference for researchers who may be looking for cost-effective, portable EEG solutions. We hope it may also serve as an inspiration for those considering incorporating portable EEG devices into their research and facilitate the conceptualisation and development of future experiments and applications.

## Acknowledgments and Funding

This review was conducted by the first author who is funded under an industry partnership grant between Macquarie University and Emotiv Pty Ltd. Emotiv contributions to this review are limited to a portion of the first author’s salary paid through Macquarie University. Emotiv did not conceive of, nor was involved in, the development of this manuscript.

**S1 PRISMA-ScR Checklist.**

**S2 Appendix. Scoping review charted data.**

